# gFACs: Filtering, Analysis, and Conversion to Unify Genome Annotations Across Alignment and Gene Prediction Frameworks

**DOI:** 10.1101/402396

**Authors:** Madison Caballero, Jill Wegrzyn

**Affiliations:** Department of Ecology and Evolutionary Biology, University of Connecticut, Storrs, CT 06269, USA

**Keywords:** Genome annotation, Bioinformatics, Protein annotation, Gene prediction, Alignment

## Abstract

Published genome annotations are filled with erroneous gene models that represent issues associated with frame, start side identification, splice sites, and related structural features. The source of these inconsistencies can often be traced to translated text file formats designed to describe long read alignments and predicted gene structures. The majority of gene prediction frameworks do not provide downstream filtering to remove problematic gene annotations, nor do they represent these annotations in a format consistent with current file standards. In addition, these frameworks lack consideration for functional attributes, such as the presence or absence of protein domains which can be used for gene model validation. To provide oversight to the increasing number of published genome annotations, we present gFACs as a software package to filter, analyze, and convert predicted gene models and alignments. gFACs operates across a wide range of alignment, analysis, and gene prediction software inputs with a flexible framework for defining gene models with reliable structural and functional attributes. gFACs supports common downstream applications, including genome browsers and generates extensive details on the filtering process, including distributions that can be visualized to further assess the proposed gene space.

## Introduction

In the era of high throughput sequencing, the size and complexity of the genomes assembled in recent years, has dramatically increased. Despite this, only a handful of the nearly 6,000 eukaryote genomes in Genbank are resolved at, or close to, chromosome level [1]. In addition, over 85% of these genomes contain some type of gene annotation errors [2-4]. These challenges are unlikely to diminish since projects, such as the Earth BioGenome Project, intend to sequence 1.5M eukaryotic genomes in coming years. Initiatives such as these will assemble increasingly large and complex genomes to assess greater biodiversity.

The majority of genome annotations are semi-automated, derived from informatic approaches that involve a combination of sequence alignments and ab initio predictions [5-8]. The inputs may include pre-assembled transcripts, raw RNA-Seq reads, and closely related proteins. The resources considered will depend on the available evidence, as well as the complexity and size of the genome under investigation. The downstream genome annotations and upstream alignment files are represented in one of the more variable bioinformatic standard file formats, known as the Generic Feature Format (GFF). The GFF file provides structure for information rich annotations as compared to the reduced representation available through GTF (General Transfer Format). Generation of a final gene annotation requires filtering of incomplete or unlikely structural models and consideration of functional annotations at the full protein or protein domain level. The informatic packages that distill several sources of evidence into gene annotations frequently deliver these without tools to assess their validity.

gFACs (gene filtering, analysis and conversion) represents a flexible annotation refining application that can accept standard annotations from primary gene annotation software as well as transcript/protein sequence aligners. This, in combination with the reference genome, can filter erroneous gene models, generate statistics/distributions, and provide outputs for standard downstream processing and/or visualization. This application does not replace the ab initio or similarity-based prediction models but serves as a companion tool to resolve conflicting annotations and improve the quality of the final models. gFACs is unique in its ability to provide statistics and analysis along with a direct connection to functional annotations to refine models. Similar programs such as gffread and gffcompare [9] provide gene model filtering and comparison abilities but lack application for analysis such as comprehensive statistics, functional annotation inclusion, and format standardizing abilities. However, gFACs can easily be used in tandem with these tools as gFACs both recognizes gffread inputs and provides a compatible GTF output for both gffread and gffcompare. The goal of gFACs is to aid the user in understanding their data while providing customized filters and utilities to remove and analyze models. As novel genomes are annotated, flexible customization and tools for analysis are essential for the tuning of final models.

## Materials and methods

Accepted inputs span a range of aligners and gene predictors, which are presented in formats with similarities to GTF and GFF files. Current accepted input formats include BRAKER/Agustus [10], MAKER [5], Prokka [11], GMAP [12], GenomeThreader[13], gffread[9], Exonerate[14], Evidence Modeler [15], and NCBI GFF annotations. The user specifies the file source at runtime, which can be selected from an applicable set of flags. gFACs will optionally accept the reference genome in FASTA format to permit more refined filtering and analysis. The second optional file, includes the annotation flat file resulting from EnTAP [16] which provides a functional annotation summary, including similarity search, protein domain, and gene family assignments for the proposed gene models or aligned sequences (**Figure 1**). The physical positions represented in these files are formatted into an intermediate text file to aid in processing and calculating the proposed gene space.

**Figure 1.**
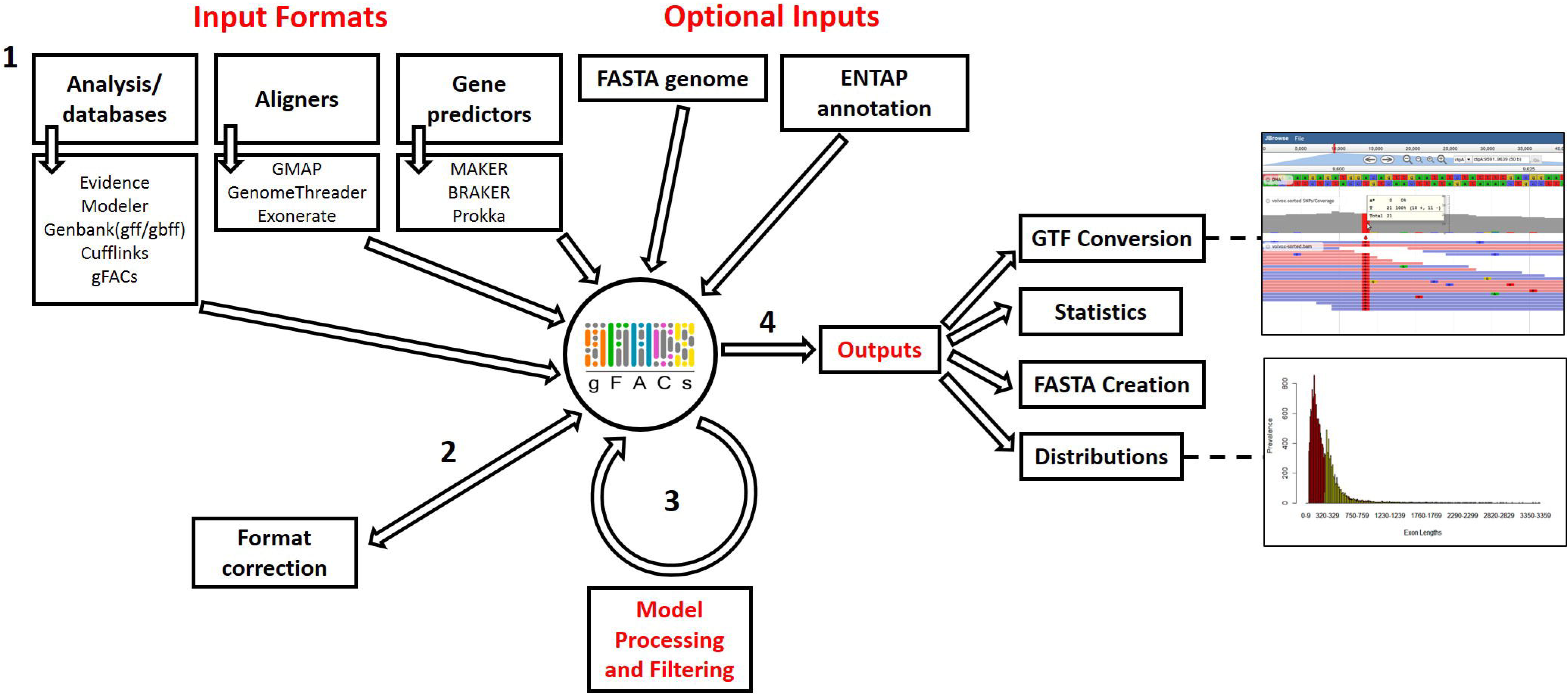
gFACs pipeline.

gFACs removes erroneous models through a set of 14 user selected filtering options (**Table 1;** Table S1), optionally aided by a reference genome or functional annotation. A notable feature in gFACs is the ability to discern and separate isoforms and conflicting models. This is performed by identifying overlapping exons due to conflicting evidence such as parent RNA. Splice site recognition is assumed from the provided annotation or alignment. Furthermore, gFACs is capable of collapsing these discerned isoforms and removing duplicate models to provide a final unique set. Each proposed model is subject to a predetermined set of filters as flagged by the user, many of which can be customized, such as setting minimum intron lengths or detection of in-frame stop codons. The addition of functional annotations will allow for the exclusion of models without a protein domain. gFACs does support alternate start codon use, which may further increase the number of passing gene models. Alternate start sites in prokaryotes are less common (up to 20% GTG or TTG) [17] and eukaryotes alternate start sites act near exclusively as rare isoforms [18]. Therefore, inclusion for alternate splice sites is user specified.

**Table 1.**
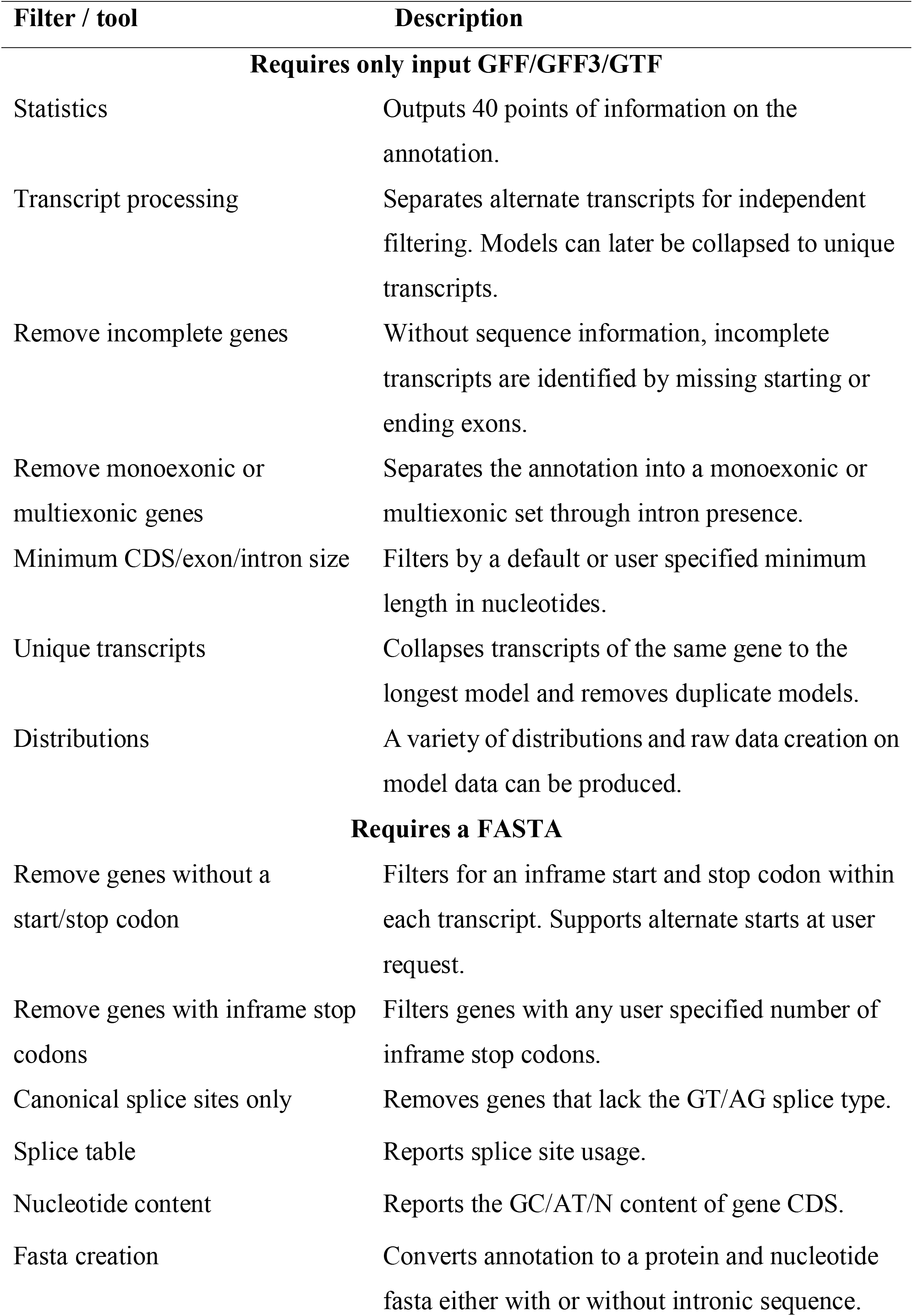
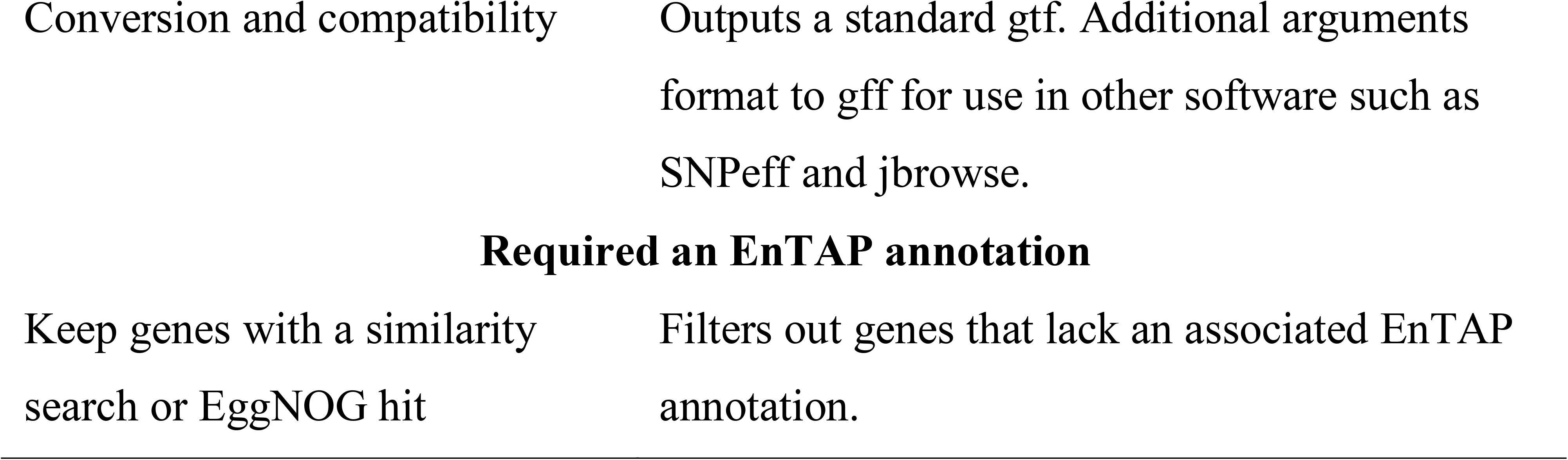
Summary of gFACs filters and utilities (simplified)

| **Filter / tool** | **Description** |
| --- | --- |
| **Requires only input GFF/GFF3/GTF** |  |
| Statistics | Outputs 40 points of information on the annotation. |
| Transcript processing | Separates alternate transcripts for independent filtering. Models can later be collapsed to unique transcripts. |
| Remove incomplete genes | Without sequence information, incomplete transcripts are identified by missing starting or ending exons. |
| Remove monoexonic or multiexonic genes | Separates the annotation into a monoexonic or multiexonic set through intron presence. |
| Minimum CDS/exon/intron size | Filters by a default or user specified minimum length in nucleotides. |
| Unique transcripts | Collapses transcripts of the same gene to the longest model and removes duplicate models. |
| Distributions | A variety of distributions and raw data creation on model data can be produced. |
| **Requires a FASTA** |  |
| Remove genes without a start/stop codon | Filters for an inframe start and stop codon within each transcript. Supports alternate starts at user request. |
| Remove genes with inframe stop codons | Filters genes with any user specified number of inframe stop codons. |
| Canonical splice sites only | Removes genes that lack the GT/AG splice type. |
| Splice table | Reports splice site usage. |
| Nucleotide content | Reports the GC/AT/N content of gene CDS. |
| Fasta creation | Converts annotation to a protein and nucleotide fasta either with or without intronic sequence. |
| Conversion and compatibility | Outputs a standard gtf. Additional arguments format to gff for use in other software such as SNPeff and jbrowse. |
| **Required an EnTAP annotation** |  |
| Keep genes with a similarity search or EggNOG hit | Filters out genes that lack an associated EnTAP annotation. |

gFACs provides a multitude of output options alongside a log detailing the gFACs process and each filtering effect. Additional options for output include gene/protein FASTA files, GTF represented models, comprehensive statistics on the selected gene models, and distribution tables and raw data of gene features. The distributions resulting from these filters can be easily imported in packages, such as R to view: gene lengths, CDS lengths, exon lengths, and exon size by order (Figure S1). Additionally, gFACs outputs can be converted for compatibility to SNPeff [19] for annotation of variants called against the genome and Jbrowse for immediate import and visualization in a web-based genome browser [20].

## Implementation

Examining protein coding gene model annotations provides insight on some of the common issues associated with gene annotations. Common problems noted in annotations include: completeness (lack of start/stop or in-frame stops), gene structure (splice sites, intron/exon lengths, and mono-exonic to multi-exonic model rations), fragmentation (incorrect start site assignment), and lack of functional validation (similarity searches, protein domains, gene family assignments). To demonstrate its utility, gFACs was applied to two public genomes (*Bos taurus*, GCF_000003055.6 [21] and *Malus domestica*, GCF_000148765.1 [22]) with the existing published annotations. Additionally, two BRAKER v2.1.0 annotations were generated here for the model *Homo sapiens* and non-model moss, *Funaria hygrometrica*. Microbial application was demonstrated with a Prokka v.1.11 annotation of the *Borrelia burgdorferi* B31, GCF_000008685.2 [10]. For human BRAKER2 predictions, RNAseq data from two libraries (Illumina 76bp PE, wgEncodeEH000146) was included with standard parameters. Similarly, *F. hydrometrica* RNAseq data is from six libraries (Illumina 150 PE, Bioproject PRJNA421369) Prokka predictions are derived from a basic run on only the genome in which candidate genes are predicted by five separate tools to provide gene candidate locations and features. All gene models (public or generated through Braker2/Prokka) were functionally annotated with EnTAP v0.8.0 utilizing the NCBI RefSeq database and EggNOG gene family database. The final models were for *H. sapiens* and *B. burgdorferi* were evaluated for precision against the public annotation following gFACs assessments.

A total of nine of the possible 14 filters were applied to the genomes, which represent unique sources: microbial, plant, and animal. These filters are: removal of all genes with an intron or exon less than 20 nucleotides, CDS size minimum of 150 nucleotides, required presence of an inframe ATG-only start and stop codon, no inframe stop codons, canonical (GT-AG) only splice sites in multiexonic genes, and an EnTAP similarity search or gene family assignment. For *B. burgdorferi*, the canonical spice filter is not used since introns do not exist. It should be noted that these filters demonstrate common issues but it would be expected that a small number of genes would have alternative splicing, micro-exons/introns, and other less common structures. These filters serve to demonstrate where there are excessive counts in these categories resulting from erroneous models.

Runtime and memory requirements vary based on the filters applied, annotation file size, and genome size. Among the species and filters described here (**Figure 2**), *B. burgdorferi* B31 (ASM868v2), with a genome size of 1.3Mb, can run on 1 CPU (2.1GHz, AMD Opteron Processor 6172) with 1GB allocated memory in 7 seconds. *M. domestica* (MalDomGD1.0; genome size: 704 Mb) ran in 9 minutes and 51 seconds on 5GB. *H. sapiens* (hg19; genome size: 3Gb), completed in 13 minutes and 46 seconds on 5GB.

**Figure 2.**
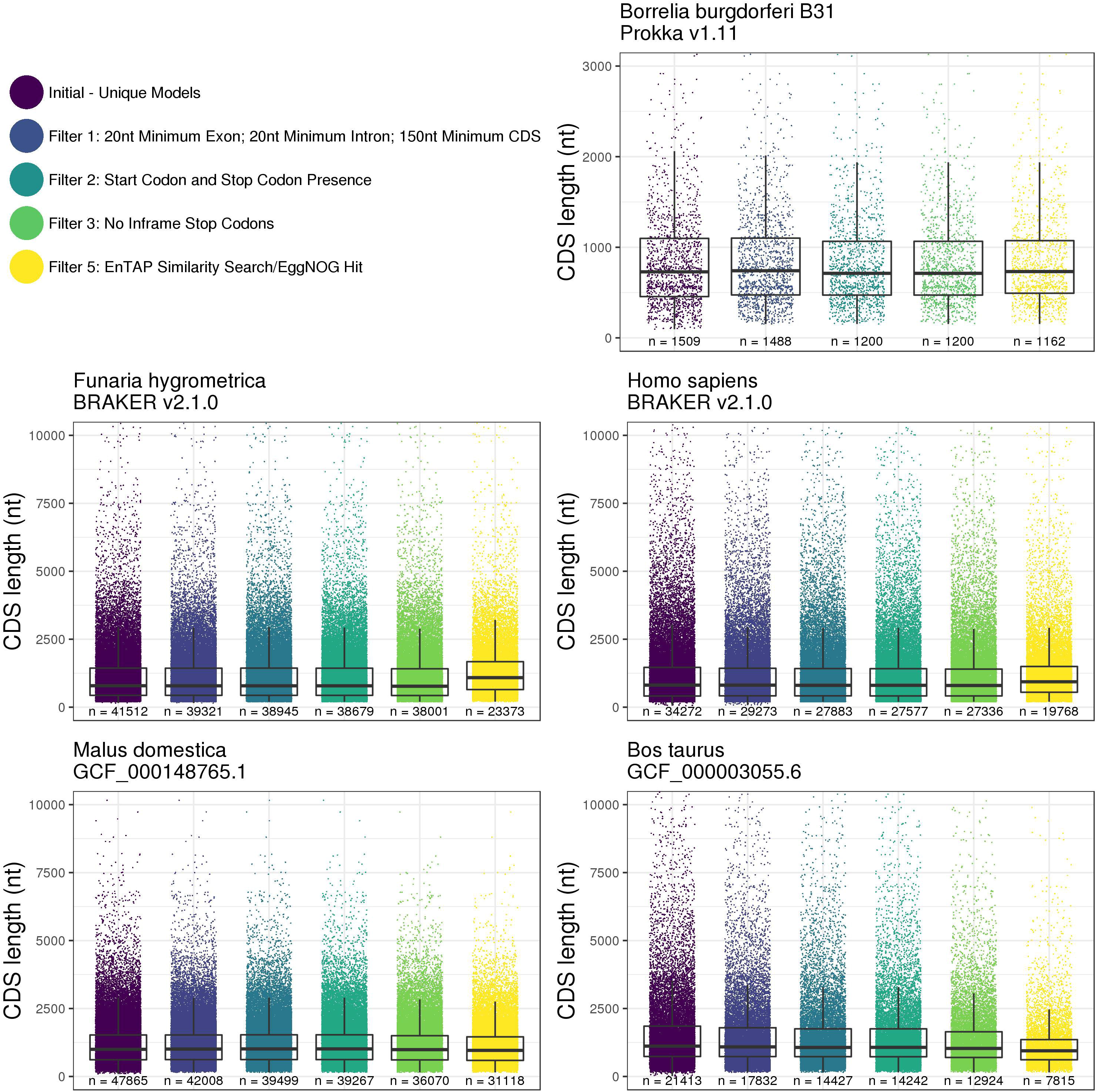
gFACs filtration on the annotations across five species reduces the number of gene models. Nine applied filters (shown as five collective filters) reduce the number of unique genes with varied intensity. Limitations on shown CDS size were placed on all species for viewing ease at 3Kb for *B. burgdorferi* (removes 85 genes) and 10 Kb of *F. hygrometrica* (168), *H. sapiens* (425), *M. domestica* (41) and *B. taurus* (262).

Across all species and annotations, gFACs was able to identify gene models that were potentially problematic (Figure 2). In *H. sapiens*, there were 34,272 unique predicted genes from BRAKER2 which was reduced to 19,768 using the applied filters (27,336 when not considering functional annotation filter). This total is comparable to the 20,203 protein coding genes of the latest human genome annotation (GRCh38p.12). Detailed analysis reveals there is a larger proportion of predicted genes that match to the same reference gene in the unfiltered annotation (22.4%) compared to the filtered annotation (12.5%). In comparing match types, the proportion of perfect matches increased from 17.0% (unfiltered) to 22.1% (filtered). The proportion of perfect and contained matches increased from 26.3% (unfiltered) to 33.8% (filtered). Finally, the proportion of all types of matches increased too with 65.5% (unfiltered) to 69.3% (filtered). To further demonstrate gFACs abilities, a more comprehensive filtering on *H. sapiens* shows stepwise reduction of models in default order and additional annotation statistics such as splice usage, nucleotide content, and 41 points of statistics on the final annotation (Table S1).

Similarly, *B. burgdorferi* predictions were reduced from 1,509 to 1,162 models which is an improvement compared to the current public annotation of 1,208 models (94.4% perfect matches, 98.9% total matches). Inclusion of alternative start codons in B. burgdorferi increases the number of passing models to 1,416 (93.2% perfect matches, 98.1% total matches) and may include a small number of erroneous models. The *F. hygrometrica, B. taurus*, and *M. domestica* models show similar rates of reductions through all filters including functional annotation at 43.70%, 34.99%, and 63.50% respectively.

The gFACs software package provides a comprehensive framework for evaluating, filtering, and analyzing gene models from a range of input applications and preparing these annotations for formal publication or downstream analysis.

## Authors’ contributions

MC developed the software package and performed all analysis in the manuscript. JW provided input into the software and helped in drafting the manuscript.

## Acknowledgements

This work was supported by the National Science Foundation Plant Genome Research Program [1444573].

We thank members of the Ecology & Evolution Biology Department at the University of Connecticut, including: Elizabeth Jockusch, Janine Caira, Hannah Ralicki, and Kaitlin Gallagher for their valuable insight and testing. We also thank the members of the Plant Computational Genomics lab at the University of Connecticut: Sumaira Zaman, Taylor Falk, and Alex Trouern-Trend for the input on the software and Nasim Rahmatpour for providing the *F. hygrometrica* annotation.

## Supplementary material

**Figure S1 Assortment of gFACs distributions rendered in R of the BRAKER2 annotations of *Pinus taeda***

Unfiltered distributions of gene feature lengths (A-D) and intron order to size (E).

**Table S1 Extensive gFACs filtering and statistics on the BRAKER 2.1.0 annotation of Homo sapiens**

and tables

Figure 1: Figure_1.jpg

Figure 2: Figure_2.jpg

Supplemental Figure 1: Figure_S1.jpg

Tables

Table 1: Table_1.docx

Supplemental Table 1: Table_S1.docx

