## Supplementary figures and images for "gFACs: Filtering, Analysis, and Conversion to Unify Genome Annotations Across Alignment and Gene Prediction Frameworks"

### Figure_S1

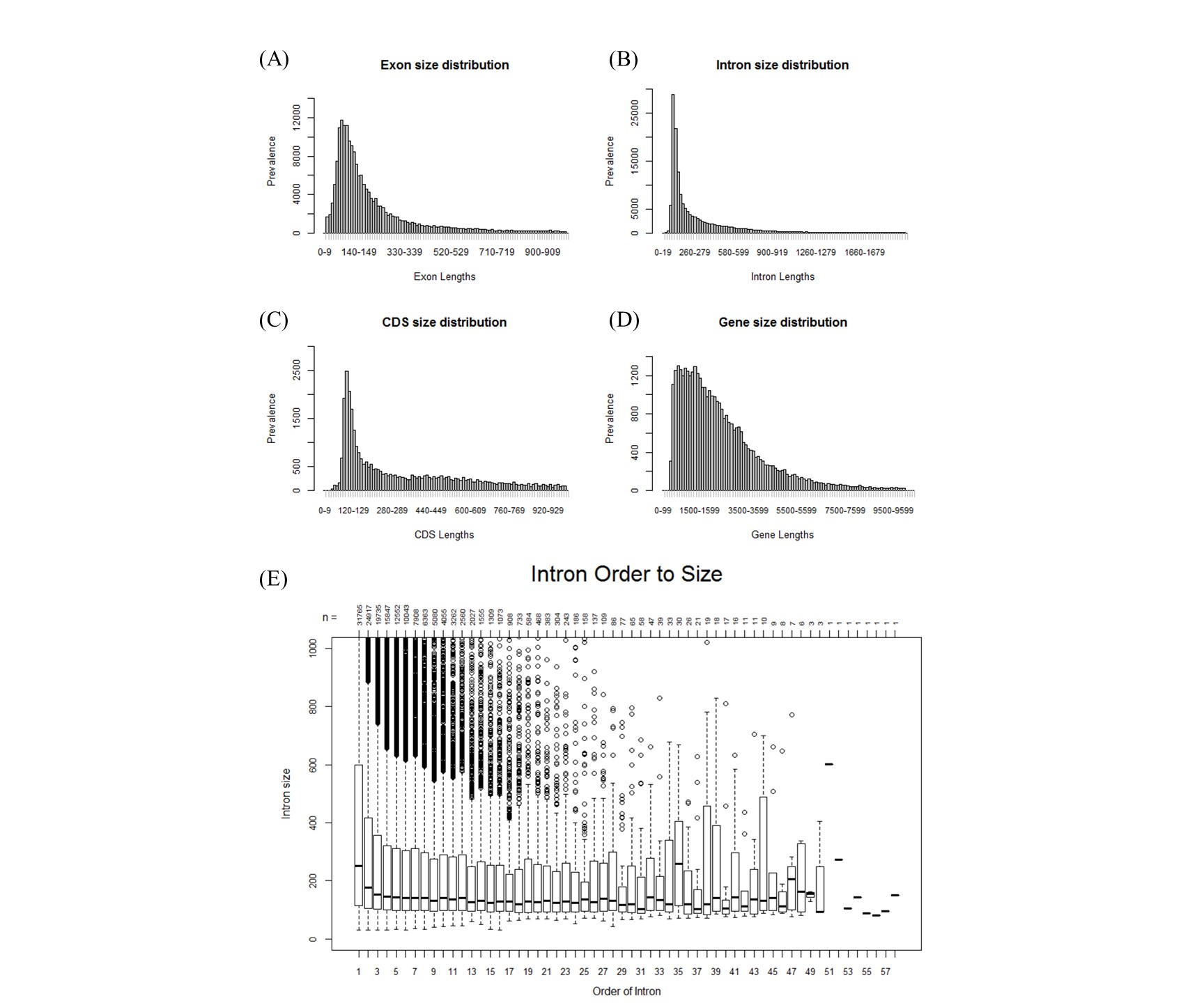
