## Supplementary material for "gFACs: Filtering, Analysis, and Conversion to Unify Genome Annotations Across Alignment and Gene Prediction Frameworks": Table_S1

**Table S1 Extensive gFACs filtering and statistics on the BRAKER 2.1.0 annotation of *Homo sapiens***

| Filter/Analysis steps | Results |
| --- | --- |
| Original input protein coding genes | 34,273 |
| Number of genes with overlap (splice/conflicting models) | 4,051 |
| Transcripts created from overlap | 9,482 |
| Number of models (overlap + nonoverlap) | 39,704 |
| Filter: Models that do not start with an intron | 38,101 |
| Filter: Models that do not end with an intron | 36,579 |
| Filter: Models that do not have an exon <20nt | 30,937 |
| Filter: Models that do not have an intron <20nt | 30,937 |
| Filter: Models that do not have a CDS <150nt | 30,937 |
| Filter: Models that have an EnTAP similarity search/ EggNOG hit | 19,768 |
| Filter: Models that have only canonical splice sites (if multiexonic) | 19,768 |
| Filter: Models that have an ATG start codon | 19,768 |
| Filter: Models that have an ending inframe stop codon | 19,768 |
| Filter: Models that have no additional inframe stop codons | 19,768 |
| Filter: Unique models (collapsed isoforms) | 19,768 |
| Analysis: Splice site type, usage count, and overall use percentage (gathered from an identical run without the canonical only filter) | gc_ag  199     0.132%   at_ac   64     0.043%  gt_ag 149957 99.83% |
| Analysis: Nucleotide content of CDS | GC content: 52.438%  AT content:   47.562%  N content: 0% |
| Statistics |  |
| Number of genes | 19,768 |
| Number of monoexonic genes | 5,761 |
| Number of multiexonic genes | 14,007 |
| Number of positive strand genes | 10,675 |
| Number of positive strand monoexonic genes | 3,394 |
| Number of positive strand multiexonic genes | 7,281 |
| Number of negative strand genes | 9,093 |
| Number of negative strand monoexonic genes | 2,367 |
| Number of negative strand multiexonic genes | 6,726 |
| Average overall gene size | 28,447.558 |
| Median overall gene size | 8,640 |
| Average overall CDS size | 1,241.413 |
| Median overall CDS size: | 933 |
| Average overall exon size | 200.118 |
| Median overall exon size | 129 |
| Average size of monoexonic genes | 766.298 |
| Median size of monoexonic genes | 660 |
| Largest monoexonic gene | 11,862 |
| Smallest monoexonic gene | 201 |
| Average size of multiexonic genes | 39,828.592 |
| Median size of multiexonic genes | 19,494 |
| Largest multiexonic gene | 1,031,336 |
| Smallest multiexonic gene | 303 |
| Average size of multiexonic CDS | 1,432.712 |
| Median size of multiexonic CDS | 1080 |
| Largest multiexonic CDS | 26,013 |
| Smallest multiexonic CDS | 201 |
| Average size of multiexonic exons | 171.715 |
| Median size of multiexonic exons | 125 |
| Average size of multiexonic introns | 5,228.523 |
| Median size of multiexonic introns | 1527 |
| Average number of exons per multiexonic gene | 8.344 |
| Median number of exons per multiexonic gene | 6 |
| Largest multiexonic exon | 14,719 |
| Smallest multiexonic exon | 20 |
| Most exons in one gene | 148 |
| Average number of introns per multiexonic gene | 7.344 |
| Median number of introns per multiexonic gene | 5 |
| Largest intron | 317,318 |
| Smallest intron | 48 |

*Note:* Filter and analysis steps are presented in the default order performed. The step of unique transcript filter (isoform collapse) is performed last by default whereas it is performed first in implementation (Figure 2). This may slightly alter the final numbers. All statistics are in nucleotide length.
